# Environmental polysulfides promote protein disulfide bond formation of microorganisms growing under anaerobic condition

**DOI:** 10.1101/2024.10.01.616119

**Authors:** Yuping Xin, Qingda Wang, Jianming Yang, Xiaohua Wu, Yongzhen Xia, Luying Xun, Huaiwei Liu

**Affiliations:** State Key Laboratory of Microbial Technology, Shandong University, Qingdao, 266200, People’s Republic of China; College of Life Sciences, Qingdao Agricultural University, Qingdao, 266109, People’s Republic of China; School of Molecular Biosciences, Washington State University, Pullman, WA, 991647520, USA

**Keywords:** Polysulfides, Protein disulfide bond, Anaerobic growth, DSB system, S-glutathionylation

## Abstract

Polysulfides are rich in anaerobic and microbial metabolism active environments. Anaerobic survival of microorganisms requires the formation of protein disulfide bond (DSB). The relation between environmental polysulfides and anaerobic DSB formation has not been studied so far. Herein, we discovered that environmental polysulfides can efficiently mediate protein DSB formation of microorganisms under anaerobic condition. We used polysulfides to treat proteins including roGFP2, Trx1, and DsbA under anaerobic condition and found that all three proteins formed intramolecular DSB *in vitro*. The growth of *E. coli* Δ*dsbB* was reduced and the amount of its intracellular protein DSB was decreased under anaerobic condition. However, treating the mutant strain with polysulfides recovered the growth and reversed DSB decrease. Treating *E. coli* Δ*dsbA* with polysulfides promoted DSB formation of its periplasmic roGFP2 and recovered its growth under anaerobic condition. In addition, treating *Schizosaccharomyces pombe* with polysulfides led to increase of the intracellular protein DSB content. Thus, our study reveals that environmental polysulfides can promote DSB formation independent of the enzymatic DSB mediating system and oxygen. In this aspect, environmental polysulfides are beneficial for the survival of microorganisms in anaerobic habitats.

**IMPORTANCE:** How polysulfides benefit adaption of microorganisms to anaerobic environments are unclear. Our study reveals that environmental polysulfides efficiently facilitate protein DSB formation under anaerobic condition. Polysulfides contain zero valent sulfur atoms (S^0^), which can be transferred to the thiol group of cysteine residue. This S^0^ atom gets two electrons from two cysteine residues and becomes reduced H_2_S, leaving two cysteine residues in disulfide bond form. Anaerobic growth of microorganisms was benefited from the formation of DSB. This finding paves the way for a deeper understanding of the intricate relationship between polysulfides and microorganisms in environmental contexts.

## INTRODUCTION

Life on earth originated in an era predating the oxygen-rich atmosphere, where primitive cells relied on sulfur respiration for energy production. In this process, hydrogen sulfide was oxidized to sulfane sulfur (S^0^) (1, 2). It is plausible to hypothesize that S^0^ played significant roles in the physiology of these early cells, and some of these roles may persist in contemporary organisms. Polysulfides, which are low molecular weight thiol compounds containing S^0^, include octasulfur (S_8_), hydrogen polysulfide (HS_n_H, n≥2), and organic polysulfide (RS_n_H, n≥2 and RS_n_R, n≥3). Polysulfides are commonly present in anaerobic habitats where microbial metabolisms are active, such as lake and coast sediments, in which their concentrations are up to 300 μM ∼400 μM (3). They are mainly produced from sulfate reduction pathway and organic sulfur (such as cysteine and methionine) degradation pathway of microorganisms. In the past two decades, polysulfides have been intensively studied and increasing evidence indicate that they are involved in a multitude of physiological processes in microorganisms, such as maintaining redox balance, regulating sulfur metabolism, and antagonizing electrophilic stress (4–6).

Disulfide bonds (DSBs) are covalent linkages that connect the sulfur atoms of two cysteine residues within a protein. Formation of DSBs is essential for both aerobic and anaerobic growth of microorganisms (7, 8). In periplasm and endoplasmic reticulum, formation of DSB is generally thought to be an enzyme-catalyzed process. Since the discovery of the first DSB-forming enzymes in 1963 (9), several key players have been identified. For example, DsbA is recognized as a pivotal enzyme in the formation of DSBs in the bacterial periplasm (10, 11). In eukaryotic cells, the endoplasmic reticulum relies on protein disulfide isomerase (PDI) (12), while oxidase Mia40 plays a central role in the mitochondria (13, 14). Under aerobic condition, these enzymatic systems use oxygen (O_2_) as the final electron acceptors. Whereas under anaerobic condition, specific electron acceptors are used. For instance, the DsbAB system uses fumarate or nitrate reductase and the PDI-Ero1 system uses flavin adenine dinucleotide (FAD) as the final electron acceptors under anaerobic condition (15–18). While these enzymatic systems are undoubtedly crucial for the formation of DSBs, they may not be absolutely essential (19, 20). It is likely that additional, yet-to-be-unveiled enzymes or factors are involved in the DSB formation process of periplasm and endoplasmic reticulum.

DSBs form also within the cytoplasm. This process is typically facilitated by low molecular weight chemical agents, such as reactive oxygen species (ROS), reactive nitrogen species (RNS), and oxidized glutathione (GSSG) (21–24). Enzymes with redox activities, such as glutaredoxin and thioredoxin, also facilitate this process. It is noteworthy that these same agents—ROS, RNS, and GSSG—also trigger S-glutathionylation (Pr-SSG), which is the formation of a mixed disulfide bond between glutathione (GSH) and a protein (25–27). Both cytoplasmic DSBs and S-glutathionylation can take place under anaerobic condition and the specific factors that mediate their formation remain to be fully elucidated.

In this study, we first performed *in vitro* experiments and found that environmental polysulfides can promote DSB formation of purified proteins under anaerobic condition. Second, we performed *in vivo* experiments and observed that environmental polysulfides can promote DSBs formation in anaerobically cultivated microorganisms. Our study revealed that environmental polysulfides are beneficial for the survival of microorganisms in anaerobic habitats via promoting DSBs formation independent of enzymatic systems.

## RESULTS

### Octasulfur mediated GSH oxidation under anaerobic condition

The small peptide GSH is prone to be oxidized into GSSG. To investigate the potential of environmental polysulfides in mediating GSH oxidation, we mixed GSH with S_8_ (5 mM of each) in KPI buffer (100 mM, pH 7.4) under anaerobic condition. After a 30-minute incubation at room temperature, the resulting products were analyzed using LC-ESI-MS. GSSG and GSSH (glutathione persulfide) were the predominant products, with MS signal intensities of 1.33×10^6^ and 1.59×10^6^, respectively (Fig. 1A, Fig. S1A, and S1B). Additionally, GSSSH and GSSSG, which are products of secondary reaction, were detected with comparatively lower MS signal intensities of 8.2×10^4^ and 2.78×10^5^, respectively (Fig. 1A, Fig. S1C, and S1D). The reaction between GSH and S_8_ was rapid. H_2_S production was detectable within 1 minute after mixing and reached a plateau within 3 minutes (Fig. 1B). No H_2_S, GSSG, GSSH, or other derivatives was detected from GSH solution without reacting with S_8_.

**FTG 1.**
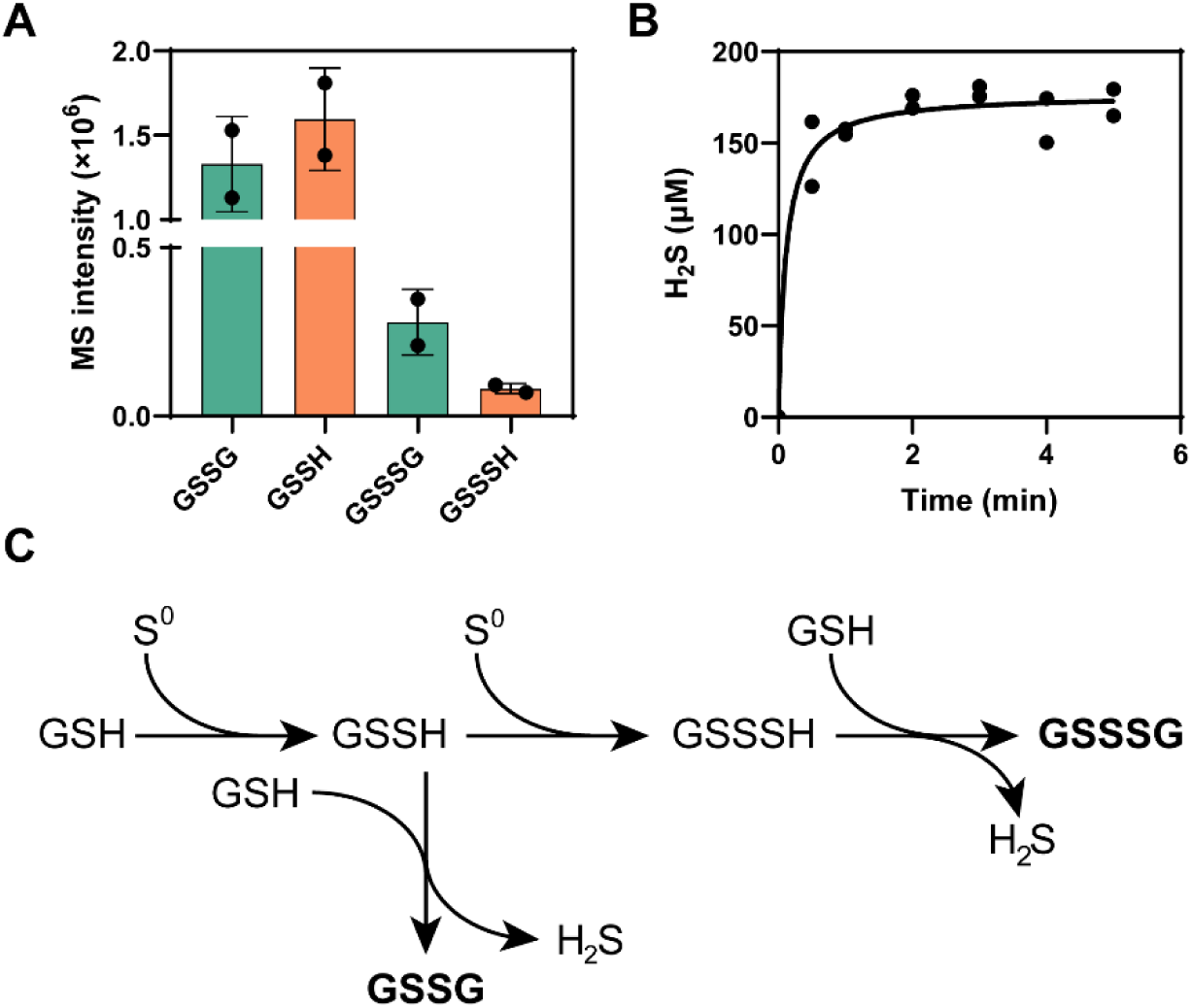
Reactions of GSH and S_8_ under anaerobic condition. (A) The MS signal intensities of GSSG, GSSH, GSSSG and GSSSH products. (B) Production of H_2_S in different reaction time. (C) The serial reactions between GSH and S_8_.

These findings suggested that environmental polysulfides can catalyze the oxidation of GSH under anaerobic condition through a series of reactions (Fig.1C). The presence of GSSH implies that S^0^ initially reacts with one GSH molecule to form GSSH. This intermediate then reacts with another GSH molecule, resulting in the formation of GSSG and H_2_S. GSSSH and GSSSG are products generated by further reactions of GSSH with S^0^ and GSH.

### Hydrogen persulfide mediated DSB formation under anaerobic condition *in vitro*

Three proteins capable of forming intramolecular DSBs were subjected to a reaction with hydrogen persulfide (HSSH) under anaerobic condition. These proteins were expressed in and purified from *E. coli* BL21(DE3), each containing an N-terminal His Tag (Fig. S2). The purified proteins were dissolved in DTT containing buffer to keep their cysteine residues reduced. The protein-HSSH reactions were conducted in KPI buffer (10 mM, pH 7.4, DTT free) at room temperature for 30 minutes under anaerobic condition.

The first protein examined was roGFP2, a widely utilized redox-sensitive fluorescent protein derived from GFP. roGFP2 features a mutation that allows its excitation spectrum to shift upon the formation of a DSB between Cys_147_ and Cys_204_, which are in close proximity to its chromophore (Fig. 2A). The formation of this DSB confers a new excitation wavelength (*E_x_*=405 nm) to roGFP2 and reduces the efficacy of the original excitation wavelength (*E_x_*=488 nm). Consequently, the ratio of emission light intensity excited at 405 nm to that excited at 488 nm (405/488) is proportional to the ratio of oxidized to reduced roGFP2 (roGFP_ox_/roGFP_re_) (28, 29).

**FIG 2.**
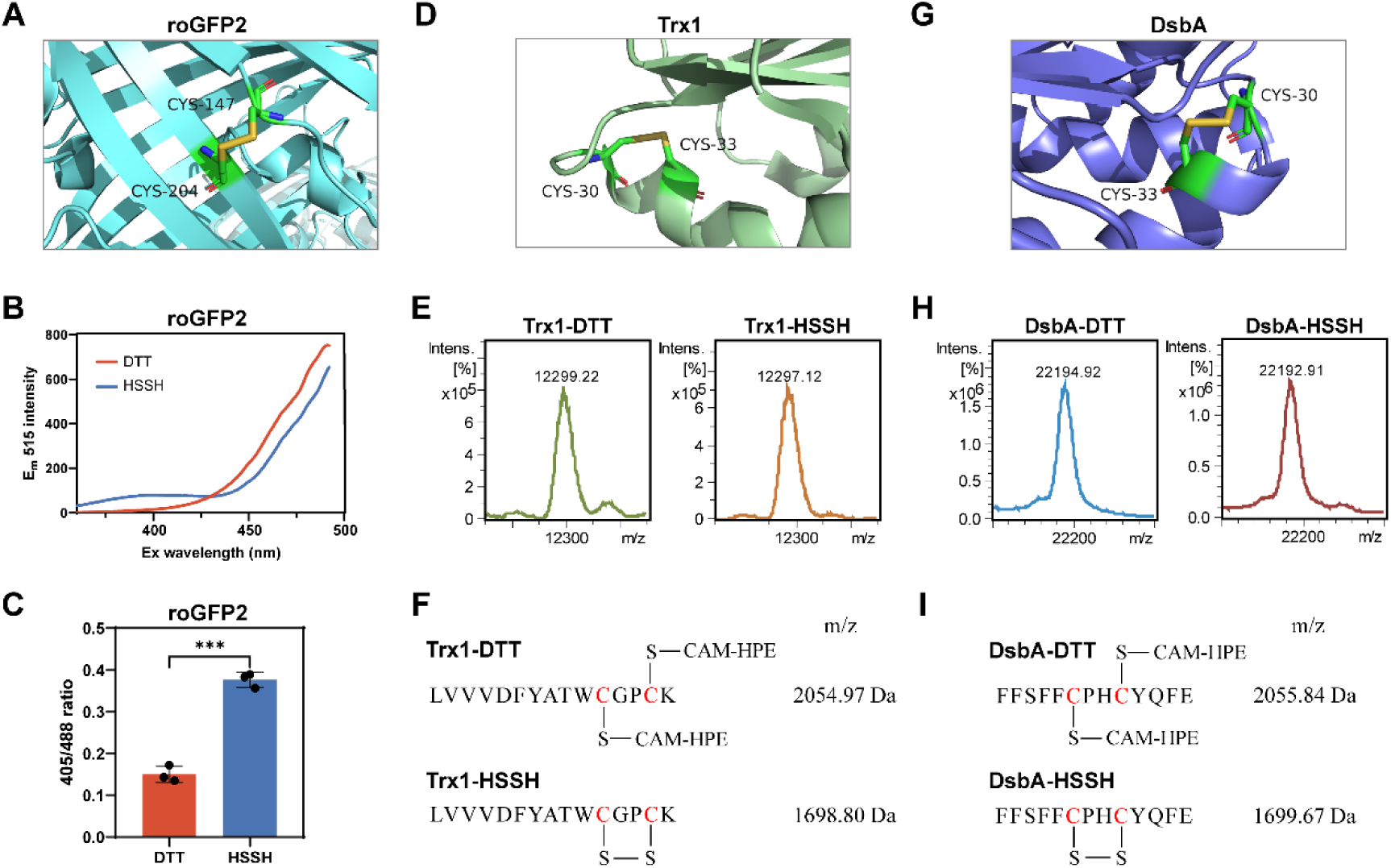
Proteins formed disulfide bonds by reacting with HSSH under anaerobic condition. (A) 3D structure of roGFP2 with a disulfide bond between Cys_147_ and Cys_204_ (PDB ID, 1JC1). (B) Full excitation spectra of roGFP2. After HSSH treatment, the excitation peak of roGFP2 around 405 nm was increased and the excitation peak around 488 nm was decreased. (C) HSSH-treated roGFP2 showed a higher ratio of 405/488 than did untreated one. (D) 3D structure of Trx1 with a disulfide bond between Cys_30_ and Cys_33_ (PDB ID, 3F3Q). (E) Full mass spectra of Trx1 and HSSH-reacted Trx1. (F) Identified peptides from untreated or HSSH-reacted Trx1. (G) 3D structure of DsbA with a disulfide bond between Cys_30_ and Cys_33_ (PDB ID, 1A2M). (H) Full protein mass spectra of DsbA and HSSH-reacted DsbA. (I) Identified peptides from untreated or HSSH-reacted DsbA. For C, data were from three independent repeats and presented as mean value ± s.d. *** represents p < 0.001 based on a two-sided t-test.

Upon examining the excitation spectrum of HSSH-reacted roGFP2, a distinct new excitation peak emerged around 405 nm, while the excitation efficacy around 488 nm was diminished (Fig. 2B). In contrast, the control, roGFP2 in DTT containing KPI buffer (denoted as DTT-reacted roGFP2), displayed only the excitation peak around 488 nm. Furthermore, the 405/488 ratio for HSSH-reacted roGFP2 was significantly higher than that of the DTT-reacted roGFP2 (Fig. 2C). These observations confirmed the formation of an intramolecular DSB in roGFP2 following its reaction with HSSH.

The second protein, thioredoxin 1 (Trx1) from *Saccharomyces cerevisiae*, is a cytoplasmic protein that participates in various redox reactions through its reversible DSB formed between Cys_30_ and Cys_33_ (Fig. 2D). Both HSSH-reacted and DTT-reacted Trx1 were analyzed using LC-MS and LC-MS/MS. At the whole protein level (analyzed by LC-MS), the molecular weight of the former was 12,297 Da, which was 2 Da less than that of the latter (12,299 Da) (Fig. 2E). At the peptide level (analyzed by LC-MS/MS), a peptide containing Cys_30_ and Cys_33_ was identified in DTT-reacted Trx1. In this peptide, Cys_30_ and Cys_33_ were found in their reduced form (-SH) and were blocked by β-(4-hydroxyphenyl)ethyl iodoacetamide (HPE-IAM) (Fig. 2F and Fig. S3). The same peptide was also identified in HSSH-reacted Trx1, but with a key difference: the cysteines formed a DSB that was not blocked by HPE-IAM (Fig. 2F and Fig. S4). These analyses confirmed the formation of a DSB in HSSH-reacted Trx1 under anaerobic condition.

The third protein, *E. coli* DsbA, is the primary facilitator of disulfide bond formation in secreted proteins. It functions by transferring its intra DSB, which is formed between Cys_30_-X-X-Cys_33_ (excluding the signal peptide), to substrate proteins (Fig. 2G). Both HSSH-reacted and DTT-reacted DsbA were subjected to LC-MS and LC-MS/MS analysis. At the whole protein MS level, the former exhibited a molecular weight of 22,192 Da, which was 2 Da lower than that of the latter (22,194 Da) (Fig. 2H). At the peptide MS level, a peptide containing Cys_30_ and Cys_33_ was detected in DTT-reacted DsbA, with a molecular weight of 2055.8 Da (Fig. 2I and Fig. S5), corresponding to the reduced form of these two cysteines (directly blocked by HPE-IAM). The same peptide was also detected in HSSH-reacted DsbA, but with a molecular weight of 1699.6 Da (Fig. 2I and Fig. S6), indicating the formation of a DSB between the two cysteine residues. Collectively, these experiments demonstrated that environmental polysulfides can mediate the formation of protein DSBs under anaerobic condition.

### Hydrogen persulfide was as effective as H_2_O_2_ in mediating DSB formation *in vitro*

To investigate the capacity of HSSH to induce DSB formation in the presence of oxygen, we conducted an experiment using 10 μM roGFP2 in various solutions: a solution containing DTT at 200 μM, one with HSSH at 200 μM, another with H_2_O_2_ at 200 μM, and a normoxic solution with dissolved oxygen at approximately 250 μM. The incubation was carried out under aerobic condition at room temperature for 30 minutes.

Analysis of the excitation spectrum revealed distinct characteristics for each solution: roGFP2 treated with HSSH exhibited a pronounced peak at 405 nm, roGFP2 treated with H_2_O_2_ showed a moderate peak; roGFP2 exposed to O_2_ had a weaker peak; and roGFP2 treated with DTT displayed no peak at 405 nm (Fig. 3A). The ratio of 405 nm to 488 nm excitation for HSSH-treated roGFP2 was significantly higher than that for DTT and O_2_ treated samples, and comparable to that for H_2_O_2_ treated one (Fig. 3B). These findings suggested that HSSH was more effective than O_2_ and as efficient as H_2_O_2_ in facilitating the formation of DSB.

**FIG 3.**
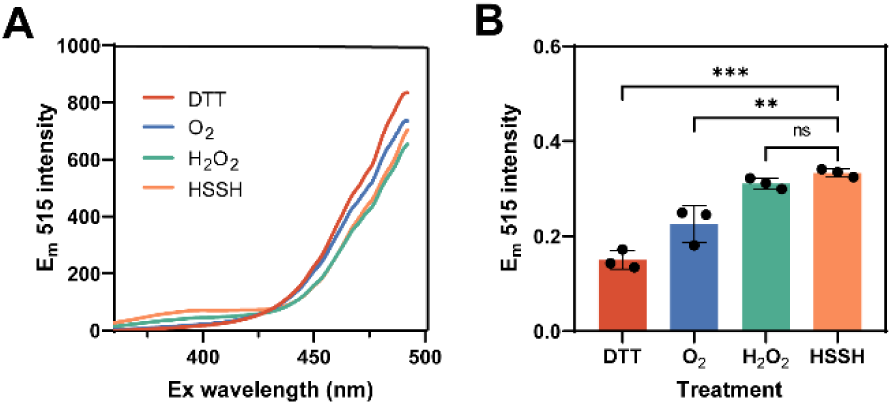
HSSH mediated roGFP2 DSB formation under aerobic condition. (A) Full excitation spectra of roGFP2. 10 μM roGFP2 was reacted with 200 μM DTT, ∼250 μM O_2_, 200 μM H_2_O_2_, or 200 μM HSSH under aerobic condition. (B) The 405/488 ratios of roGFP2. Data were from three independent repeats and presented as mean value ± s.d. ** represents p < 0.01 and *** represents p < 0.001 based on a two-sided t-test.

### Octasulfur compensated for DsbB deletion in *E. coli*

The DsbB protein serves as the oxidizing agent for DsbA, and together they mediate DSBs formation in periplasm of *E. coli*. We constructed a strain of *E. coli* MG1655 with *dsbB* deletion, termed Δ*dsbB*, and firstly cultivated it in LB medium under aerobic condition. Compared to the wild-type *E. coli* MG1655 strain (wt), the Δ*dsbB* strain exhibited a subtle yet significant growth impairment (Fig. 4A), indicating that disrupting DSB system leads to growth defects. We added 0.2 mM S_8_ into Δ*dsbB* cultivation medium and found that, interestingly, growth of Δ*dsbB* was partially recovered.

**FIG 4.**
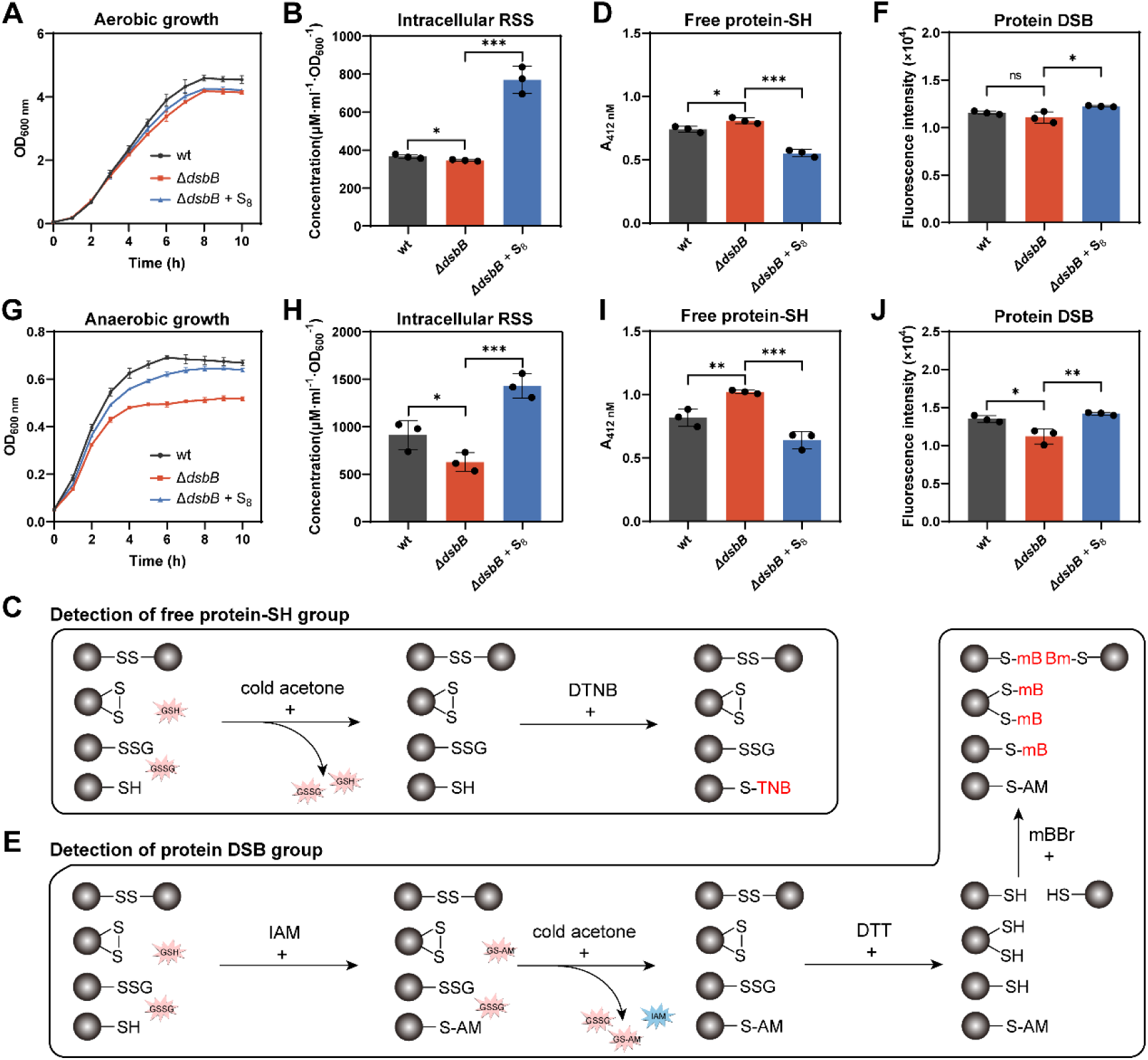
S_8_ complemented *dsbB* knock-out under both aerobic and anaerobic conditions. (A) Growth of *E. coli* MG1655 wt, Δ*dsbB*, and S_8_-treated Δ*dsbB* under aerobic condition. (B) Amounts of intracellular polysulfides in *E. coli* MG1655 strains under aerobic condition. (C) Workflow of the DTNB based free protein-SH group quantification. (D) Amounts of the total free protein-SH group in *E. coli* MG1655 strains under aerobic condition. (E) Workflow of the mBBr based protein DSB group quantification. (F) Amounts of the total protein DSB group in *E. coli* MG1655 strains under aerobic condition. (G) Growth of *E. coli* MG1655 strains under anaerobic condition. (H) Amounts of intracellular polysulfides in *E. coli* MG1655 strains under anaerobic condition. (I) Amounts of the total free protein-SH group in *E. coli* MG1655 strains under anaerobic condition. (J) Amounts of the total protein DSB group in *E. coli* MG1655 strains under anaerobic condition. Data were from three independent repeats and presented as mean value ± s.d. * represents p < 0.05, ** represents p < 0.01, and *** represents p < 0.001 based on a two-sided t-test.

To further confirm the effect of S_8_, we collected 50 mL *E. coli* cells (OD_600_=1, cultivated under aerobic condition) and resuspended them in 25 mL PBS buffer with or without 0.5 mM S_8_. After an incubation at room temperature for 30 min, cells were collected and subjected to further analysis. Without S_8_ treatment, the Δ*dsbB* strain contained slightly fewer intracellular polysulfides than the wt strain. When Δ*dsbB* strain was treated with S_8_, its intracellular polysulfides significantly increased and became about 2-fold higher than that of wt (Fig.4B). Subsequently, the intracellular total free protein thiol (-SH) groups were quantified. Cells were harvested and then lysed using a high-pressure homogenizer. Their proteomes were isolated from low molecular weight molecules (including GSH and its oxidized form GSSG) using cold acetone precipitation (Fig. 4C). A probe that selectively tags free -SH groups, 5,5’-Dithiobis (2-nitrobenzoic acid) (DTNB), was used to label the proteomes. After reaction with DTNB, the absorbance at 412 nm was measured. The Δ*dsbB* proteome displayed a higher absorbance value than did the wt proteome, indicating that the portion of proteins in -SH status was higher in Δ*dsbB* than that in wt (Fig. 4D). When Δ*dsbB* was treated with S_8_, the absorbance of its proteome became lower than that of wt proteome, suggesting that S_8_ decreased the portion of proteins in -SH status.

We then quantified the intracellular total DSB groups of the cells. After cell disruption, free protein-SH groups were blocked with iodoacetamide (IAM), and the proteomes were again isolated using cold acetone precipitation. The proteomes were then treated with DTT to convert their DSBs into free protein-SH groups. The resulting DSB-derived protein-SH groups were labeled with monobromobimane (mBBr) and quantified by measuring fluorescence intensity (Fig. 4E). There was no significant difference in fluorescence intensity between the Δ*dsbB* and wt proteomes (Fig. 4F). However, when Δ*dsbB* was treated with S_8_, the fluorescence intensity of its proteome significantly increased, suggesting that S_8_ increased the portion of proteins in DSB status.

Secondly, we cultivated Δ*dsbB* in LB medium under anaerobic condition and examined its growth again. Different from results obtained from aerobic condition, the Δ*dsbB* strain showed significantly impaired growth compared to wt strain under anaerobic condition (Fig. 4G), but the growth was significantly recovered by adding 0.2 mM S_8_ into cultivation medium. These observation indicated that under anaerobic condition, environmental polysulfides were more important for *E. coli* cells compared to what they were under aerobic condition.

Furthermore, we collected 100 mL *E. coli* cells (OD_600_=0.5, cultivated under anaerobic condition) and resuspended them in 25 mL PBS buffer (deoxygenated by purging with nitrogen for 30 minutes) with or without 0.5 mM S_8_. After an incubation at room temperature for 30 min under anaerobic condition, cells were collected and subjected to further analysis. When Δ*dsbB* was treated with S_8_, its intracellular polysulfides increased about 3-fold (Fig. 4H). The Δ*dsbB* strain had a substantially higher concentration of free protein-SH groups and a significantly lower concentration of protein DSB groups than the wt strain (Fig. 4I and 4J). S_8_ treatment significantly decreased the amount of free protein-SH groups and increased the protein DSB amount of Δ*dsbB* proteome.

In summary, these findings underscore the critical role of DsbB in DSB formation, but in the absence of DsbB, treating cells with environmental polysulfides can compensate for its function, with the compensatory effect being particularly pronounced under anaerobic condition.

### Environmental polysulfides compensated for DsbA deletion in *E. coli*

DsbA directly reacts with target proteins to promote the formation of their DSBs. We also constructed a strain of *E. coli* MG1655 with *dsbA* deletion, termed Δ*dsbA*. As Δ*dsbB,* Δ*dsbA* showed also significant growth impairment under anaerobic condition and S_8_ treatment recovered the growth (Fig. 5A). To visually detect the influence of *dsbA* deletion on DSBs formation of periplasm, we expressed the roGFP2 protein and located it in periplasm of both wt and Δ*dsbA* strains. The strains were then treated with 0.5 mM S_8_, HSSH, GSSG, DTT, or H_2_O_2_ under anaerobic condition. We observed that the ratio of oxidized roGFP2 (OxD) was quickly increased by S_8_ and HSSH treatment in both wt and Δ*dsbA* strains (Fig. 5B and 5C). GSSG and H_2_O_2_ also increased OxD, but their increasing rates were slower than that of S_8_ and HSSH. In aspect to the reoxidation rate calculated from the data of first 10 minutes treatment, no obvious difference was observed between wt and Δ*dsbA* (Fig. 5D). These results suggested environment polysulfides can promote periplasmic DSB formation independent of DsbA, and the promoting effects were more efficient than GSSG or H_2_O_2_.

**FIG 5.**
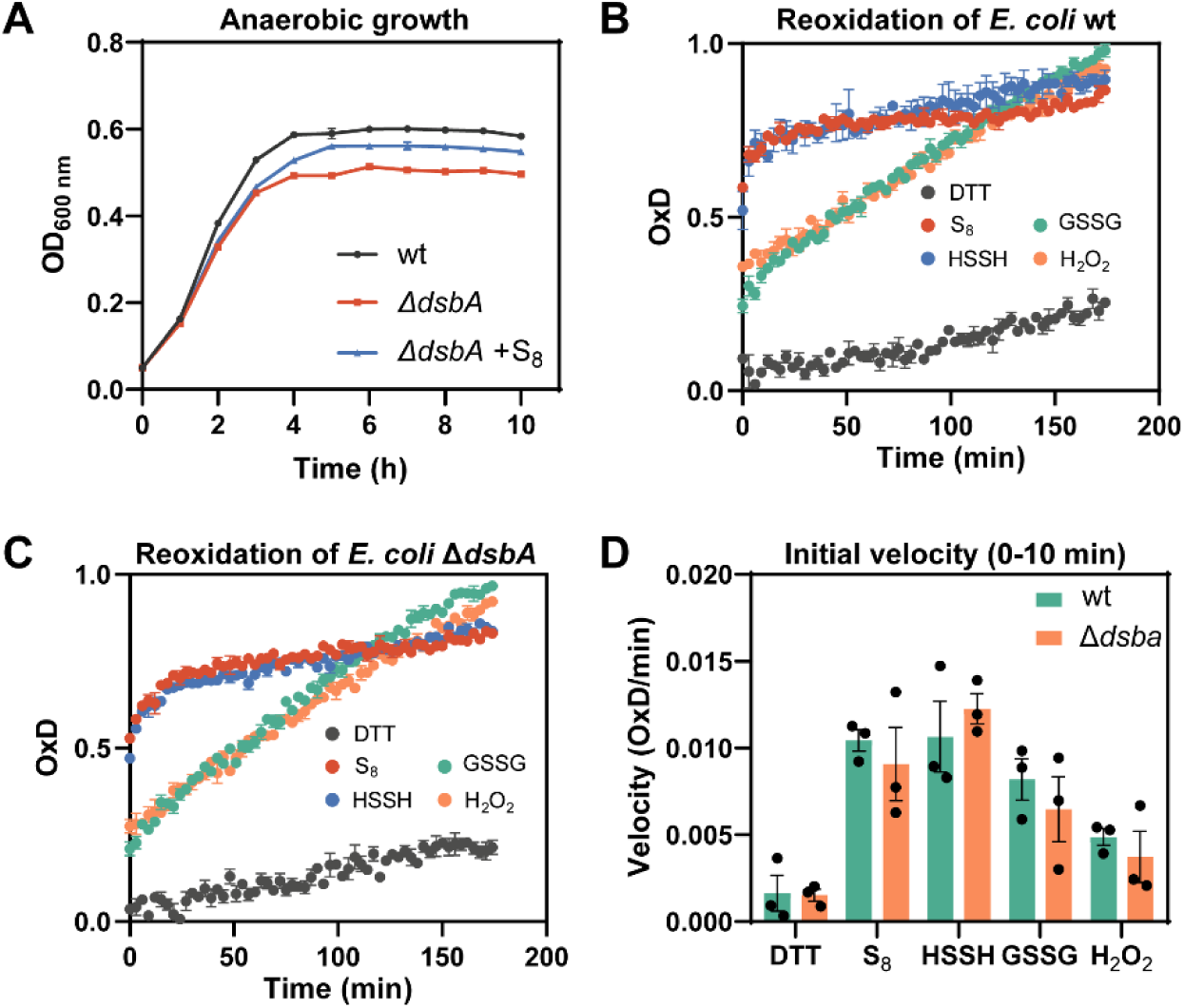
Environmental polysulfides compensated periplasmic roGFP2 re-oxidation velocity in *E. coli* lacking DsbA. (A) Growth of *E. coli* MG1655 wt, Δ*dsbA*, and S_8_-treated Δ*dsbA* under aerobic condition. (B, C) Effect of various sulfur-containing compounds on the reoxidative capacity of roGFP2 in the periplasmic space of *E. coli*. E. *coli* was reduced with 10 mM DTT, washed to remove excess DTT, and treated with 0.5 mM S_8_, HSSH, GSSG, or H_2_O_2_. DTT-reduced cells served as a control. The fluorescence changes of roGFP2 were monitored over 180 minutes. (D) The initial reoxidation rate of roGFP2 in *E. coli* wt and Δ*dsbA* was measured during the first 10 minutes of the assay. Data were from three independent repeats and presented as mean value ± s.d.

### GSSH mediated protein S-glutathionylation

To assess the ability of GSSH in facilitating protein S-glutathionylation, which is the inter DSB between proteins and GSH, we first mixed GSSH with cysteine at a 1:1 molar ratio under anaerobic condition. The reaction proceeded at room temperature for 30 minutes, and the products were subsequently analyzed using LC-ESI-MS. For comparative purpose, we also performed a reaction using GSSG as a positive control under identical conditions, as GSSG is known to mediate protein S-glutathionylation. The product cysteine-SSG was detected in both GSSH and GSSG reaction mixtures, with the MS signal intensities being 2.6×10^6^ for GSSH and 4.6×10^6^ for GSSG (Fig. 6A and 6B). These results indicated that both GSSH and GSSG can react with –SH group to form S-glutathionylation.

**FIG 6.**
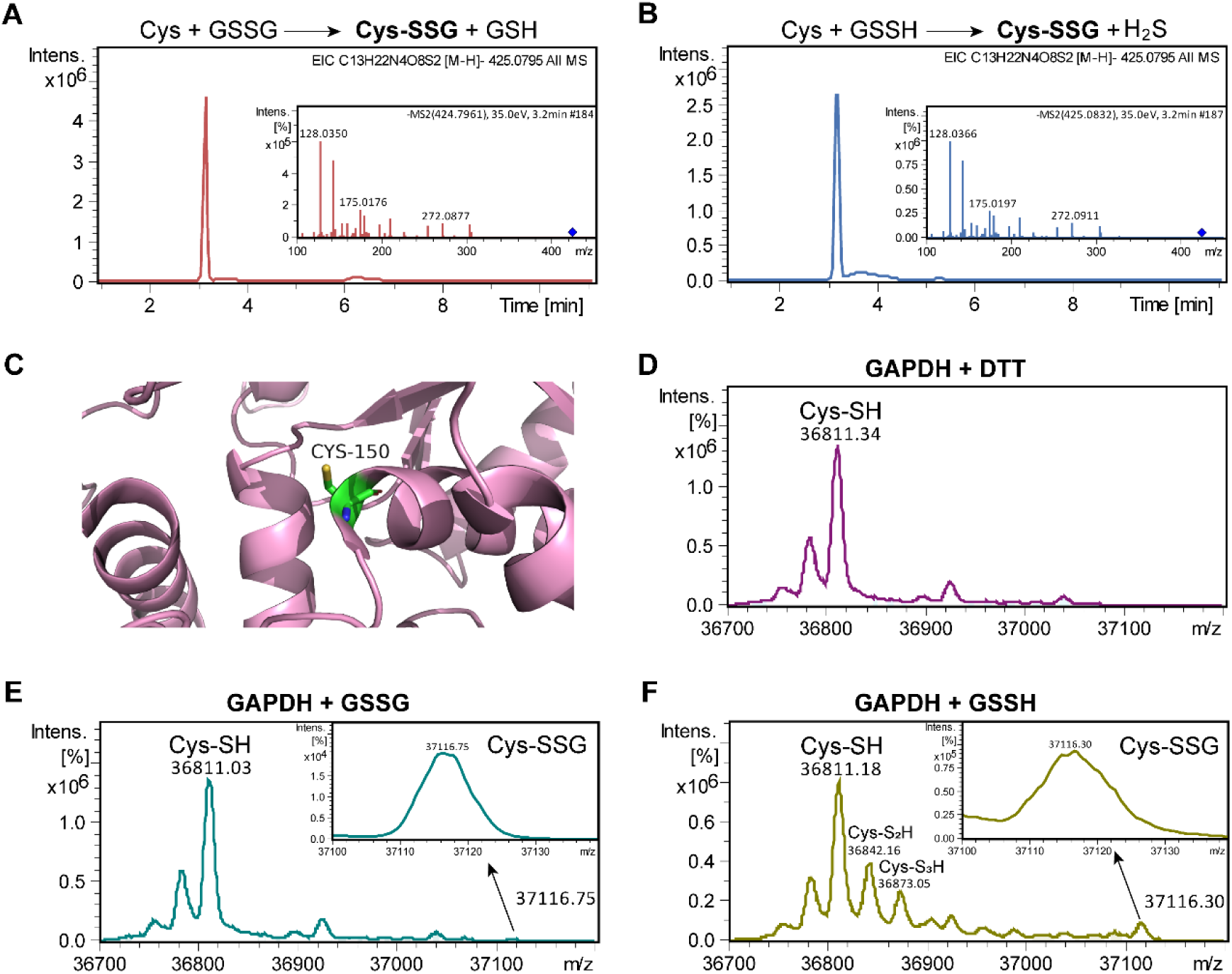
GSSH mediated S-glutathionylation formation in GAPDH. (A) LC-ESI-MS analysis of products from GSSG (5 mM) and cysteine (5 mM) reaction. (B) LC-ESI-MS analysis of products from GSSH (5 mM) and cysteine (5 mM) reaction. (C) 3D structure of GAPDH with Cys_150_ (PDB ID, 3PYM). (D) Full protein mass spectra of DTT-reacted GAPDH. (E) Full protein mass spectra of GSSG-reacted GAPDH. (F) Full protein mass spectra of GSSH-reacted GAPDH.

*S. cerevisiae* glyceraldehyde 3-phosphate dehydrogenase (GAPDH) is known to undergo S-glutathionylation in response to oxidative stress (Fig. 6C) (30). We procured purified GAPDH and mixed it with an excess of DTT, GSSG, or GSSH under anaerobic condition. After a 1-hour incubation at room temperature, the modified GAPDH was analyzed by LC-MS. The DTT treated GAPDH displayed a principal molecular weight peak of 36,811 Da, which matched its calculated molecular weight of reduced form (Fig. 6D). For GSSG and GSSH treated GAPDH, a minor molecular weight peak of 37,116 Da appeared, which was corresponding to the GAPDH-SSG complex (Fig. 6E and 6F). Notably, the signal intensity of GAPDH-SSG was approximately 4-fold higher in GSSH treated GAPDH sample than that in GSSG treated sample. These findings indicate that, similar to GSSG, GSSH can also mediate GAPDH S-glutathionylation.

It is noteworthy that except for GAPDH-SSG, we also detected GAPDH-S_2_H and GAPDH-S_3_H from GSSH reacted sample, but these modifications were not detected from GSSG reacted sample, indicating that GSSH can also lead to protein polysulfidation modification while GSSG has no such activity.

### Octasulfur increased DSB amount in eukaryotic microorganism

To explore the influence of environmental polysulfides on DSB formation of eukaryotic cells, we treated *S. pombe* cells (OD_600_=2) with 0.5 mM S_8_ under anaerobic condition. Following a 30-minute incubation at room temperature, the intracellular polysulfide content was measured. Exposure to S_8_ resulted in a substantial increase in intracellular polysulfides, approximately a 6-fold enhancement relative to the untreated control group (Fig. 7A). Concurrently, there was a notable reduction in the free protein thiol (-SH) groups and a corresponding increase in protein DSB groups within the S_8_-treated cells (Fig. 7B and 7C).

**FIG 7.**
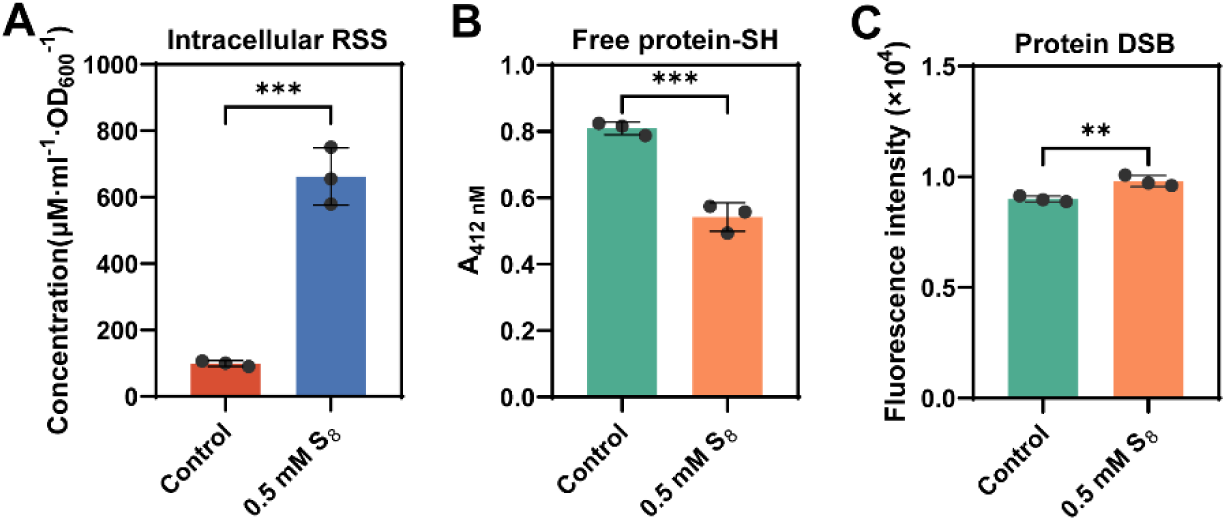
Correlation of intracellular polysulfides level with protein DSB level in eukaryotic cells *S. pombe*. (A) Treating *S. pombe* with S_8_ increased intracellular polysulfides level. (B) Treating *S. pombe* with S_8_ caused decrease in total free protein-SH group level. (C) Treating *S. pombe* with S_8_ caused increase in total protein DSB group level. Data were from three independent repeats and presented as mean value ± s.d. * represents p < 0.05 and *** represents p < 0.001 based on a two-sided t-test.

## DISCUSSION

In this study, we first verified that environmental polysulfides can oxidize GSH to GSSG under anaerobic condition. Building on this, we treated proteins, including roGFP2, Trx1, and DsbA, with polysulfides under anaerobic condition and observed the formation of intra-molecular DSB in all three proteins. Further investigation involved the deletion of DsbB in *E. coli*. We observed a significant decrease in protein DSB level in the mutant strain. Notably, treatment with S_8_ was able to reverse this decrease, with the most pronounced effects observed under anaerobic condition. Treatment with S_8_ and HSSH promoted DSB formation of periplasmic roGFP2 quickly in DsbA deletion strain. Additionally, treating eukaryotic cells resulted in increase of intracellular DSB content. Collectively, these results suggested that under anaerobic condition, environmental polysulfides can promote DSB formation in dependent of DsbAB system.

Recent work by Knoke et al. (31) reported that in DsbA-deficient *E. coli*, roGFP2 located in the periplasm can still form DSB, indicating the presence of an alternative system or player for DSB formation. They proposed that GSSG might directly oxidize periplasmic roGFP2 in DsbA-deficient *E. coli*. However, previous studies, including their own, showed that direct oxidation of roGFP2 by GSSG was inefficient. They speculated that an unidentified, GSH-dependent factor could oxidize roGFP2 in the absence of DsbA. Our previous studies have shown that intracellular polysulfides content was influenced by GSH levels, and vice versa (32, 33). We have now demonstrated that environmental polysulfides can directly oxidize roGFP2, providing an answer to their speculation—the unidentified factor is likely intracellular polysulfides.

The redox potentials of the *E. coli* periplasm and cytoplasm are approximately −160 mV and −260 mV, respectively (34, 35). DsbA has a high redox potential value of −120 mV among all known thiol–disulfide oxidoreductases (36), enabling it to maintain an oxidized state and mediate protein DSB formation in the periplasm. The standard redox potential of sulfane sulfur (S^0^ + 2H^+^ + 2e^−^ → H_2_S) is +144 mV, indicating that it can oxidize most proteins in both the periplasm and cytoplasm.

When S_8_ reacts with GSH, GSSH is produced as an intermediate, suggesting a two-step reaction mechanism (Reactions 1 and 2).

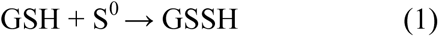

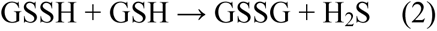

We hypothesize that polysulfides oxidize proteins through a similar mechanism. Initially, polysulfides transfer a sulfane sulfur atom to a cysteine residue on the target protein, forming a protein-SSH intermediate. Subsequently, the protein-SSH reacts with another cysteine to form a DSB, while the sulfane sulfur is reduced to H_2_S.

A previous report showed that GSSH can react with the glucose-induced biofilm accessory protein A (GbaA) to form GbaA-SSG (37). Herein, we found that GSSH can also react with GAPDH to form GAPDH-SSG. This activity of GSSH should be ascribed to its thiol-oxidizing activity conferred by the sulfane sulfur atom. The reaction between GSSH and proteins to form protein S-glutathionylation likely follows the same mechanism as reaction (2). In addition to DSB and S-glutathionylation, polysulfides can mediate a third modification, protein polysulfidation (protein-S_n_H, n≥2). Actually, when using GSSH to react with GAPDH, we observed the generation of GAPDH-S_2_H and GAPDH-S_3_H (Fig. 5F), which were not observed from GSSG-reacted GAPDH. Therefore, the intracellular total free protein thiol groups (-SH) quantified with our method should contain a portion of protein-S_n_H groups. However, due to lack of reliable protein-S_n_H quantification method, their contents cannot be determined as so far (38, 39). On the other hand, the intracellular DSB groups quantified with our method may modestly affected by protein-S_n_H groups, because a recent report indicated that protein-S_n_H groups labeled with IAM (protein-S_n_-AM) were not stable and easily converted to protein-S-AM (40), which should not disturb DSB quantification.

It is important to note that treating DsbA deletion strain with environmental polysulfides can quickly oxidize its periplasmic roGFP2. For DsbB deletion strain, the compensatory effect of S_8_ also was more pronounced under anaerobic condition compared to aerobic condition. This correlates with the observation that polysulfides commonly accumulate in anaerobic habitats, where DsbAB system cannot work efficiently and oxygen related species are scarce. Historically, polysulfides predated oxygen and reactive oxygen species (ROS). In the early stages of life, polysulfides may have already been mediating DSB formation. With the advent of the oxygen-rich era, this function persisted, becoming necessary primarily under anaerobic condition.

## MATERIALS AND MATHODS

### Strains and materials

All strains, human cells, and plasmids are listed in Supplementary Table S1. *Escherichia coli* strains were grown in lysogeny broth (LB) medium at a temperature of 37°C with shaking (200 rpm). The construction of the *dsbB* knockout strain was carried out according to a previously published method (41). *S. pombe* HL6381 strains, characterized by the genotype (*h^+^ his3-D1 leu1-32 ura4-D18 ade6-M210*), were cultivated in yeast extract medium (YES) at 30°C with shaking (200 rpm).

GSH, GSSG, S_8_ (99.9% purity), sodium hydrosulfide (NaHS), cysteine, DTNB, and DTT were procured from Sigma-Aldrich (Shanghai, China). S_8_ solution was prepared as reported previously (42). Briefly, a saturation solution was made by dissolving excess sulfur powder in acetone. The concentration of saturated acetone sulfur is determined as 17 mM. HSSH was prepared following the established protocol (43). GSSH was synthesized using the method outlined in (44).

### Chemical reactions of polysulfides with thiols

For reaction of S_8_ with GSH, reacting solutions used were purged with nitrogen for 30 minutes to remove dissolved oxygen, and then were placed in the anaerobic incubator. The mixing and incubating processes were conducted in the anaerobic incubator. GSH (5 mM) were prepared in KPI buffer (100 mM, pH 7.4). S_8_ solution was prepared by dissolving S_8_ powder in methanol in sealed bottle. GSH solution was mixed with S_8_ (5 mM) at room temperature. For the reactions of cysteine with GSSG or GSSH, the experiments also were conducted in the anaerobic incubator with nitrogen purged solutions. GSSG and GSSH solutions were prepared in KPI buffer (100 mM, pH 7.4). Cysteine (5 mM) was mixed with GSSG and GSSH solutions (5 mM) in an anaerobic incubator. Following a 30-minute incubation period at room temperature.

For analysis, the reacted mixtures were subjected to centrifugation at 12,000 *g* for 3 minutes. The resulting supernatant was subsequently analyzed using liquid chromatography-electrospray ionization mass spectrometry (LC-ESI-MS).

### LC-ESI-MS analysis

The analysis procedure was performed as previously described (39). In summary, the samples underwent liquid chromatography (LC) using an InertSustain C18 column (Shimadzu, Japan) interfaced with a High-Resolution Q-TOF mass spectrometer (Ultimate 3000, Bruker impact HD, Germany). The samples were introduced into the LC system via the loading pump, with 0.25% acetic acid serving as the mobile phase A and 100% methanol as the mobile phase B. Over the initial minute, the concentration of mobile phase B was ramped up from 7.5% to 52.5% and held constant for 15 minutes. At the 15-minute mark, the concentration of mobile phase B was further increased to 55%, then to 100%, and sustained for an additional 5 minutes. At 20.1 minutes, mobile phase B was reduced to 7.5% and maintained at this level until the completion of the analysis at 31 minutes. The electrospray ionization (ESI) source temperature was set to 200°C, and the ion spray voltage was set at 4.5 kV. Nitrogen was employed as the nebulizer and drying gas. The acquired data were processed using Data Analysis 4.2 software.

### Protein expression and purification

*E. coli* BL21(DE3) strains containing the expression plasmids pET30-roGFP2, pET30-Trx1, pET30-DsbA, or pET30-Gapdh were cultivated in LB medium supplemented with 100 μg/mL kanamycin. Upon reaching an optical density (OD_600_) of 0.6–0.8, isopropyl β-D-1-thiogalactopyranoside (IPTG) was introduced to a final concentration of 0.3 mM to induce protein expression, and the culture was continued for an additional 18 hours at 20°C. The bacterial cells were harvested by centrifugation and then re-suspended in a lysis buffer (50 mM NaH_2_PO_4_, 300 mM NaCl, 20 mM imidazole, 0.5 mM DTT, pH 8.0). Cell lysis was achieved using a high-pressure homogenizer, model SPCH-18 (Stansted), and the lysate was clarified by centrifugation at 12,000 g for 15 minutes. The supernatant was subsequently applied to a Ni-NTA agarose affinity resin for protein purification, following the manufacturer’s protocol. The purified protein was desalted using a PD-10 desalting column (GE Healthcare) that had been pre-equilibrated with a desalination buffer (0.5 mM DTT, 50 mM Tris-HCl, 10% glycerin, pH 7.4). The finally obtained protein solutions were stored in icebox in the anaerobic incubator before reacting with HSSH and other reagents. The purity of the protein was assessed by sodium dodecyl sulfate-polyacrylamide gel electrophoresis (SDS–PAGE), and protein concentration was quantified using a BCA protein assay kit (Beyotime Biotechnology, China).

### Polysulfides and ROS reactions with proteins

For the reaction of roGFP2 with HSSH, H_2_O_2_, dissolved O_2_, and DTT under aerobic condition. The dissolved oxygen concentration was measured using a dissolved oxygen (DO) probe. The purified roGFP2 was diluted to a concentration of 0.3 mg/mL (10 μM) in KPI buffer (10 mM, pH 7.4). Subsequently, 100 μM of HSSH, H_2_O_2_, or DTT was introduced to the roGFP2 solution. The mixtures were incubated at room temperature for 30 minutes.

For the reaction of roGFP2 with HSSH under anaerobic condition, reacting solutions used were purged with nitrogen for 30 minutes to remove dissolved oxygen. The mixing, incubating, and desalting processes were all conducted in the anaerobic incubator. The roGFP2 solution (10 μM) was mixed with 100 μM HSSH in KPI buffer (10 mM, pH 7.4) and incubated at room temperature for 30 minutes. Following the incubation period, the mixtures were processed through a PD-10 desalting column to eliminate any unreacted low molecular weight compounds. The roGFP2 samples post-reaction were then prepared for fluorescence analysis. The excitation spectra of roGFP2 were recorded using an RF-5301 PC fluorescence spectrophotometer (Shimadzu, Japan). The fluorescence intensities emitted at 511 nm (*E_m_*=511 nm), when excited at 405 nm (*E_x_*=405 nm) and 488 nm (*E_x_*=488 nm), were quantified using a Synergy H1 microplate reader (BioTek, USA). The ratio of the emitted light intensities at these two excitation wavelengths (405/488) was calculated to assess the redox state of roGFP2.

For the reaction of Trx1 and DsbA with HSSH under anaerobic condition, reacting solutions used were purged with nitrogen for 30 minutes to remove dissolved oxygen. The mixing, incubating, and desalting processes were all conducted in the anaerobic incubator. Trx1 and DsbA were prepared at a concentration of 1.8 mg/ml (150 μM) and 3.2 mg/ml (150 μM), respectively. These were diluted in KPI buffer (10mM, pH 7.4). Subsequently, 1.5 mM of HSSH was introduced into the mixture, which was then incubated under anaerobic conditions at room temperature for a duration of 30 minutes. Following the incubation period, the mixture underwent purification through a PD-10 desalting column to eliminate any unreacted low molecular weight reagents. The obtained protein samples were subsequently analyzed using LC-MS and LC-MS/MS.

For the reactions of GAPDH with GSSG and GSSH under anaerobic condition, reacting solutions used were purged with nitrogen for 30 minutes to remove dissolved oxygen. The mixing, incubating, and desalting processes were all conducted in the anaerobic incubator. GAPDH was diluted to 1.1 mg/ml (30 μM) in KPI buffer (10 mM, pH 7.4), 300 μM of GSSG or GSSH was added and the mixture was incubated at room temperature for 1 h. After the incubation, the mixture was passed through a PD-10 desalting column to remove unreacted low molecular weight reagents. The obtained proteins were subjected to LC-MS analysis.

### Protein LC-MS and LC-MS/MS analysis

The molecular weight of whole protein was analyzed using LC-MS. LC system equipped with an XBridge Protein BEH C4 Sentry Guard Cartridge (Waters™, USA) and High-Resolution Q-TOF mass spectrometry (Ultimate 3000, Burker impact HD, GER) were used. The mobile phase A consisted of 0.1% formic acid (FA), while mobile phase B was a mixture of acetonitrile and 0.1% formic acid (FA). Initially, the concentration of mobile phase B was set at 5% for the first 7 minutes of the analysis. At the 7-minute mark, the mobile phase composition was dynamically adjusted to a blend comprising 10% of 0.1% FA and 90% of the acetonitrile/0.1% FA mixture. This mixture was maintained for a duration of 3 minutes, after which mobile phase B was reduced back to 5% and this condition was sustained for an additional 3 minutes. The flow rate throughout the process was meticulously controlled at 0.5 ml/min. The Q-TOF mass spectrometer was equipped with an electrospray ionization (ESI) source operating in positive ion mode, with a capillary voltage set to 3500 volts. The acquired data were subjected to deconvolution and comprehensive analysis using the Data Analysis 4.2 software.

For LC-MS/MS analysis, protein sample was treated with HPE-IAM and subsequently digested with trypsin (0.5 mg/ml). The generated peptides were desalted by using a C18 column, eluted in 70% acetonitrile and 0.1% trifluoroacetic acid and freeze-dried. The final product was resuspended in HPLC grade water. LC-MS/MS analysis was performed using the Prominence nano-LC system (Shimadzu, Shanghai, China) equipped with a custom-made silica column (75 μm×15 cm) packed with 3 μm Reprosil-Pur 120 C18-AQ. The elution process involved a 100-minutes gradient ranging from 0% to 100% of solvent B (0.1% formic acid in 98% acetonitrile) at a flow rate of 300 nl/min. Solvent A was composed of 0.1% formic acid in 2% acetonitrile. The eluent was ionized and electrosprayed via the LTQ-Orbitrap Velos Pro CID mass spectrometer (Thermo Scientific, Shanghai, China), which operated in data-dependent acquisition mode using Xcalibur 2.2.0 software (Thermo Scientific). Full-scan MS spectra (ranging from 400 to 1800 m/z) were detected in the Orbitrap with a resolution of 60,000 at 400 m/z.

### Growth analysis of *E. coli* with S_8_ treatment

*E. coli* strains were initially cultivated in LB broth overnight to ensure exponential growth. Subsequently, these cultures were subcultured into fresh LB medium (OD_600_= 0.05), either supplemented with or without S_8_ solution (the final concentration of S_8_ in medium was 0.2 mM). The cultivation was conducted under both aerobic and anaerobic conditions. For the aerobic condition, 300 mL-scale shake-flasks containing 100 mL LB and inoculated *E. coli* were placed in a 37°C incubator with shaking (200 rpm). For the anaerobic condition, the sealed serum bottles (250 mL scale) containing 100 mL LB medium and inoculated *E. coli* were purged with nitrogen for 30 minutes to remove dissolved oxygen. Samples were taken out using a 1 mL-scale syringe. The incubation temperature was maintained at 37°C throughout the experiment. The growth curves were determined by measuring the optical density of OD_600_ at 1-hour intervals over a period of 10 hours.

### S_8_ treatment of E. coli and Schizosaccharomyces pombe

*E. coli* strains was cultivated in 50 ml LB medium at 37°C with shaking (200 rpm) under aerobic or anaerobic condition. For *E.coli* cultivated under aerobic condition, 50 ml cells were collected when OD_600_ reached 1. Cells were diluted in 25 mL PBS buffer (10 mM, pH 7.4) to make its OD_600_=2, then 0.7 mL S_8_ solution was added (the final S_8_ concentration was 0.5 mM). After 30 min incubation at 37°C with shaking (200 rpm), cells were collected and subjected to intracellular polysulfides, protein free thiol groups, and protein disulfide bonds analysis. For *E. coli* cultivated under anaerobic condition, 100 mL cells were collected when OD_600_ reached 0.5. The following processes were performed in the anaerobic incubator: cells were diluted in 25 mL PBS buffer (10 mM, pH 7.4, purged with nitrogen for 30 minutes before using) to make its OD_600_=2, then 0.7 mL S_8_ solution was added (the final S_8_ concentration was 0.5 mM). After 30 min incubation at 37°C with shaking (200 rpm), cells were collected and resuspened in different reaction buffers suitable for intracellular polysulfides, protein free thiol groups, and protein disulfide bonds analysis.

*S. pombe* HL6381 was cultivated in 50 mL YES medium at 30°C with shaking (200 rpm). When OD_600_ reached 1, cell cultures were collected. Cells were diluted in 25 mL PBS buffer (10 mM, pH 7.4) to make its OD_600_=2, then 0.7 mL S_8_ solution was added (the final S_8_ concentration was 0.5 mM). After 30 min incubation at 30°C with shaking (200 rpm), cells were collected and subjected to different reaction buffers suitable for intracellular polysulfides, protein free thiol groups, and protein disulfide bonds analysis.

### Quantification of intracellular polysulfides in *E. coli* and *S. pombe*

For *E. coli* and *S. pombe* cells, S_8_ treatment was performed as described above. The intracellular levels of polysulfides were quantified as a method previously reported in the literature (45). Briefly, 1.5 mL S_8_ treated *E. coli* and *S. pombe* cells obtained from one culture dish were collected and resuspended in 100 uL reaction buffer (50 mM Tris-HCl, 1% Triton X-100, 50 mM sulfite, and 50 μM diethylene triamine pentaacetic acid, pH 9.5). This suspension was subjected to thermal treatment by incubating at 95°C for 10 minutes, a step designed to convert intracellular polysulfides into thiosulfate. Following the incubation, the reaction mixture was centrifuged to separate the components. A 50 μL aliquot of the supernatant was carefully pipetted out for further analysis. This supernatant was then incubated with 5 μL of monobromobimane (mBBr) solution at a concentration of 25 mM, at room temperature for 30 minutes to facilitate the derivatization of thiosulfate. To terminate the reaction, 55 μL of a mixture of acetic acid and acetonitrile (in a volume ratio of 1:9) was added. After the addition of the stopping solution, the mixture was centrifuged to collect the supernatant layer. The concentration of thiosulfate present in this supernatant was determined using high-performance liquid chromatography (HPLC) on a Shimadzu system from Japan.

### Detection of intracellular protein free thiol groups and protein disulfide bonds

For *E. coli* and *S. pombe*, S_8_ treatment was performed as described above. For detection of intracellular protein free thiol groups, 25 mL S_8_ treated *E. coli* and *S. pombe* cells (OD_600_=50) were collected and resuspended in 10 mL PBS buffer (10 mM, pH 7.4) supplemented with 1% protease inhibitors to prevent unwanted proteolysis. Subsequently, the cell suspensions were subjected to lysis using a high-pressure homogenizer, specifically the SPCH-18 model from Stansted. The lysed cells were then centrifuged at 12,000 *g* for 15 minutes to separate the soluble proteins in the supernatant. The supernatant was treated with cold acetone to precipitate the proteins. A volume of 500 μL of the supernatant was mixed with 1.5 mL of 100% pre-chilled acetone and the mixture was incubated at −20°C for 90 minutes to facilitate protein precipitation. Following the incubation, the mixture was centrifuged again at 14,000 *g* for 10 minutes to pellet the proteins. The supernatant was carefully removed, and the acetone was allowed to evaporate at room temperature, yielding a protein pellet. The protein pellet was then resuspended in a denaturing buffer (8 M urea) to solubilize the proteins. The resulting protein solution was diluted to a concentration of 5 mg/mL. To this solution, 20 mM DTNB was added to initiate the reaction. This mixture was incubated at room temperature for 15 minutes. Finally, the light absorbance of the reacted proteins at a wavelength of 412 nm was measured using a Synergy H1 microplate reader from BioTek, USA.

The intracellular concentration of protein DSB was quantified as previously reported (46). Briefly, 25 mL S_8_ treated *E. coli* and *S. pombe* cells (OD_600_=50) were collected and resuspended in 10 mL PBS buffer solution (100 mM, pH 8.0) supplemented with 20 mM NaCl, 1 mM EDTA, 200 mM iodoacetamide (IAM), and 1% protease inhibitor. The cell suspensions were then subjected to lysis using the high-pressure homogenizer SPCH-18 from Stansted. Following lysis, the supernatant was harvested by centrifugation at 12,000 *g* for 15 minutes. A 500 μL aliquot of the supernatant was treated with IAM and incubated at 37°C for 1 hour in a light-protected environment to alkylate and block any free thiol groups present in the proteins. To eliminate unreacted IAM, the protein solution was precipitated using a three-fold volume of pre-chilled acetone and incubated at −20°C for 90 minutes. After this incubation, the mixture was centrifuged at 14,000 *g* for 10 minutes. The supernatant was carefully removed, and the acetone was allowed to evaporate at room temperature, yielding a protein pellet. The protein pellet was then resuspended in a denaturing buffer (8 M urea). The protein solution was diluted to a concentration of 1 mg/ml and mixed with 1 mM dithiothreitol (DTT). This mixture was incubated at 37°C for 30 minutes to reduce any disulfide bonds present in the proteins.

Subsequently, the newly liberated thiol groups were reacted with 2.5 mM monobromobimane (mBBr), and the reaction was allowed to proceed at room temperature for 20 minutes. The fluorescence resulting from the mBBr-thiol adduct was measured using a Synergy H1 microplate reader from BioTek, USA. The fluorescence intensities were recorded with excitation at 370 nm (*E_x_*=370 nm) and emission at 485 nm (*E_m_*=485 nm).

### Detection of roGFP2 reoxidation capacity in the periplasmic space of *E. coli*

To localize roGFP2 into the periplasmic space, plasmids were constructed following the method described in (47). PCR was performed to amplify the torA signal sequence using the MG1655 genome as a template. The torA signal sequence was fused with sequence of roGFP2 and the resulting product was then ligated into the plasmid ptrc99a using In-Fusion assembly master mix (Takara, JP).

The reoxidative capacity of roGFP2 in *E. coli* wt and Δ*dsbA* strains was assayed based on the method described in (31). Plasmid ptrc99a-torA(SP)-roGFP2 was first transformed into *E. coli* wt and Δ*dsbA* strains. These strains were cultured in LB medium at 37°C and 200 rpm until the OD_600_ reached ∼0.4-0.6. Then, 0.5 mM IPTG was added, and the cultures were incubated at 20°C for 16 hours. The cells were collected and washed twice with HEPES buffer (40 mM, pH 7.4, purged with nitrogen for 30 minutes to remove dissolved oxygen), resuspended in the same buffer to adjust the OD_600_ to 1.0 under anaerobic condition. The resuspended cells were mixed with 10 mM DTT or 1 mM H_2_O_2_, and incubated at room temperature for 30 minutes. To detect the reoxidative capacity of different reagents, cells were first treated with 10 mM DTT to reduce the periplasmic roGFP2, and then were washed with deoxygenated HEPES buffer to remove excess DTT. The reagents including 0.5 mM S_8_, 0.5 mM HSSH, 0.5 mM GSSG, 0.5 mM H_2_O_2_, or 0.5 mM DTT were then used to treat these cells under anaerobic condition. After 30 minutes incubation at room temperature, 100 µL of the bacterial suspension was transferred to a wells of a black, clear-bottom 96-well plate, and fluorescence was detected using a Synergy H1 microplate reader (BioTek, USA) with emission at 515 nm and excitation at 405 nm and 488 nm, respectively. The oxidation state of the probe was calculated using the fluorescence intensity ratio at the excitation wavelengths of 405 and 488 nm. All values were normalized using the following equation:

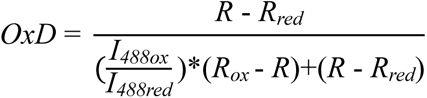

where *R_ox_* and *R_red_* are the 405 nm/488 nm ratios for full oxidized (1 mM H_2_O_2_ treatment) and full reduced (10 mM DTT treatment) roGFP2, respectively. *I_488ox_* and *I_488red_* are the fluorescence intensities of roGFP2 at 488 nm under oxidizing and reducing conditions, respectively. *R* is the 405 nm/488 nm ratio of roGFP2 in *E. coli* strains treated with different reagents.

## Statistical Analysis

Statistical analysis was done using GraphPad Prism 9.0. Data are presented as mean ± S.D. The significant difference between the two groups was analyzed using an independent student’s t test; the p-value < 0.05 indicated statistical significance.

## Data and materials availability

All data needed to evaluate the conclusions in the paper are present in the paper and/or the Supplementary Materials. Additional data related to this paper may be requested from the authors.

## ACKNOWLEDGMENTS

We thank Jing Zhu and Yuyu Guo from the Analysis and Testing Center of the State Key Laboratory for Microbial Technology (Shandong University) for assistance with the LC-MS/MS analysis.

## AUTHOR CONTRIBUTIONS

H.L.: conceptualization, funding acquisition, supervision. L.X.: conceptualization, supervision. Y.X.: investigation. Q.W. and J.Y. Methodology. X.W.: resources and data curation. Y.X.: Validation. All authors have given approval to the final version of the manuscript.

## SUPPLEMENTARY MATERIALS

**Fig S1.** Mass spectra of GSSH, GSSG, GSSSH, and GSSSG that were produced from S_8_ + GSH reaction.

**Fig S2.** SDS-PAGE analysis of roGFP2, Trx1, DsbA, Gapdh.

**Fig S3.** LC-MS/MS spectra showing that intramolecular disulfide bond formed between C_30_ and C_33_ in the LVVVDFYATWCGPCK peptide of Trx1.

**Fig S4.** LC-MS/MS spectra showing that C_30_ and C_33_ in FFSFFCPHCYQFE peptide of DsbA was directly blocked by HPE-IAM.

**Fig S5.** LC-MS/MS spectra showing that intramolecular disulfide bond formed between C_30_ and C_33_ in FFSFFCPHCYQFE peptide of DsbA.

**Fig S6.** LC-MS/MS spectra showing that C_30_ and C_33_ in LVVVDFYATWCGPCK peptide of Trx1 were directly blocked by HPE-IAM.

**Table S1.** Strains and plasmids used in this study.

**Table S2.** Primers used in this study.

